# Sex-specific long-term alteration of hippocampal excitation/inhibition balance and behavior by transient caffeine exposure during synaptogenesis

**DOI:** 10.64898/2026.08.28.747753

**Authors:** Solen Rimbert, Jessica Pressey, Ferran Gomez-Castro, Zahra Imani, Marion Russeau, David Blum, Marika Nosten-Bertrand, Sabine Lévi

**Affiliations:** ESPCI, CNRS UMR 8249, PSL Université, 75005 Paris, France; INSERM UMR-S 1270, Sorbonne Université, Institut du Fer à Moulin, 75005, Paris, France; UMR-S1172 Lille Neuroscience & Cognition (LilNCog), University of Lille, Inserm, CHU Lille, F-59000 Lille, France; NeuroSU - site du Fer à Moulin, UMR CNRS 8265, INSERM 1341, 75005 Paris

**Author notes:** Correspondence to: **Sabine Lévi,** Brain Plasticity Lab CNRS UMR-8249, 10 rue Vauquelin, 75005 Paris.

## Abstract

Caffeine is the most widely consumed psychoactive substance worldwide, yet the long-term consequences of exposure during critical periods of brain development remain incompletely understood. Synaptogenesis represents a vulnerable window during which environmental factors can shape the maturation of neuronal circuits and influence lifelong brain function. Here, we investigated the impact of caffeine exposure during hippocampal synaptogenesis on synaptic development, neuronal function, behavior, and seizure susceptibility, with a particular focus on sex-dependent effects.

Developmental caffeine exposure induced distinct, sex-specific trajectories of hippocampal synaptic remodeling. In the CA1 region, caffeine produced opposite patterns of glutamatergic synapse regulation, characterized by a delayed reduction in excitatory synapse density in males and an increase in females. In contrast, inhibitory synapse organization was selectively altered in males, with a transient increase in CA3 inhibitory synaptic density during development associated with enhanced inhibitory transmission, whereas females exhibited no significant changes. These findings reveal sex-specific and temporally divergent effects of developmental caffeine exposure on hippocampal synaptic maturation and function.

At the behavioral level, developmental caffeine exposure produced distinct sex-dependent phenotypes : males exhibited increased anxiety-like behavior, whereas females developed a delayed impairment in recognition memory that became apparent only in adulthood. Furthermore, caffeine exposure selectively increased PTZ-induced seizure susceptibility in juvenile females, an effect that was no longer detected in adulthood.

Together, these findings demonstrate that caffeine exposure during hippocampal synaptogenesis induces sex-specific and temporally dynamic alterations in circuit maturation, resulting in distinct behavioral and neuronal excitability outcomes. These results highlight the importance of considering both sex and developmental timing when assessing the neurodevelopmental consequences of caffeine exposure.

**Graphical abstract:** 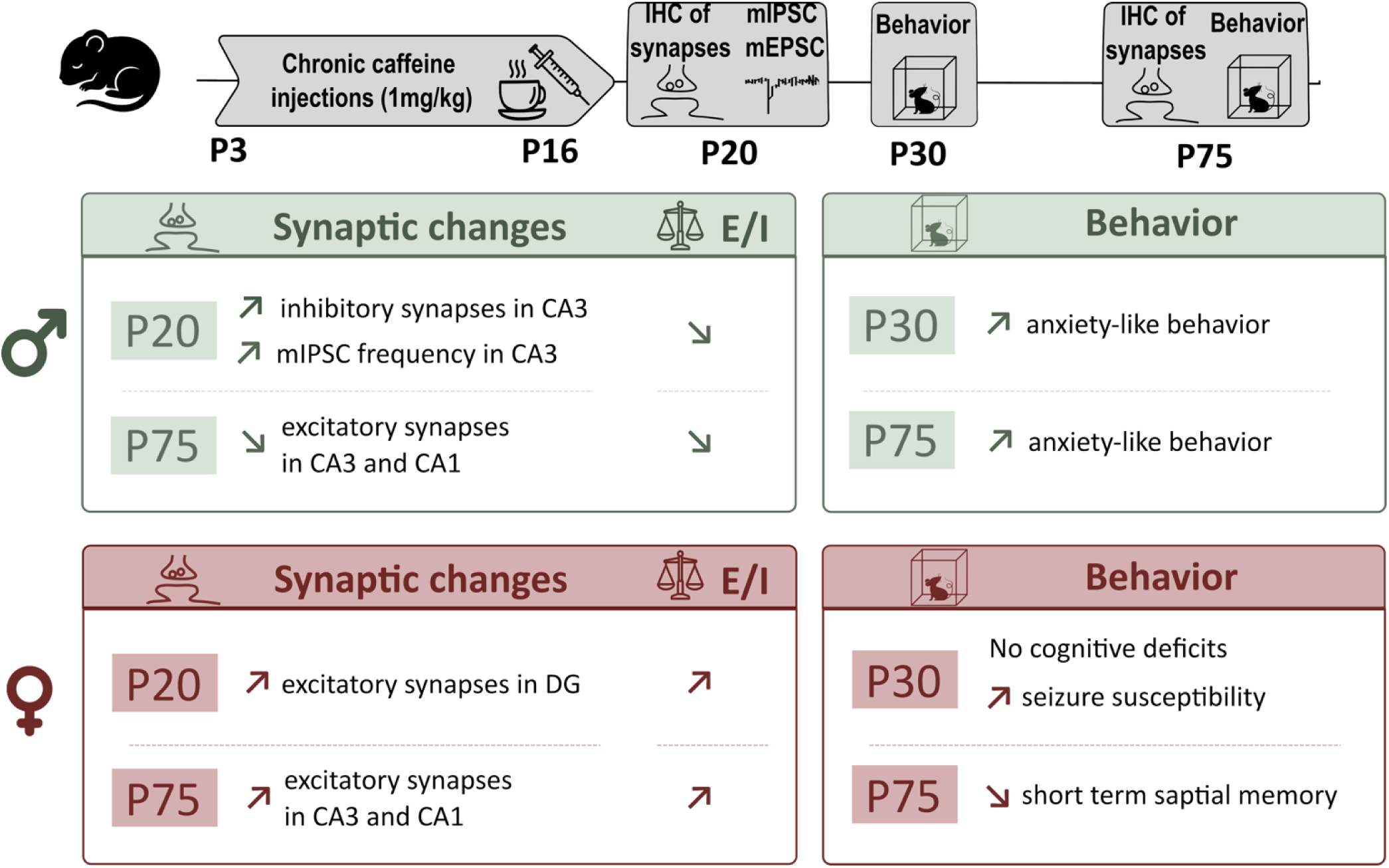

## Introduction

Caffeine is the most widely consumed psychoactive substance worldwide. It acts primarily as a competitive antagonist of adenosine A₁ and A₂A receptors (A_1_R and A_2A_R), two G protein-coupled receptors that regulate neuronal excitability, synaptic transmission, and plasticity [1]. Because caffeine readily crosses the blood–brain barrier, plasma concentrations achieved after moderate daily consumption are sufficient to block central A_1_R and A_2A_R. Caffeine is also commonly consumed during pregnancy and lactation, raising concerns that early-life exposure may interfere with endogenous adenosine signaling during critical stages of brain maturation.

Prenatal and early postnatal caffeine exposure impairs the migration of hippocampal GABAergic interneurons, resulting in hippocampal hyperexcitability, increased seizure susceptibility, and cognitive deficits [2]. Similar alterations have been reported in the cerebral cortex, where developmental caffeine exposure delays interneuron migration and integration, leading to enhanced network activity [3]. Therefore, transient disruption of adenosine signaling during development can produce persistent alterations in neuronal circuit assembly.

Increasing evidence suggests that developmental caffeine exposure affects synaptic maturation in a sex-dependent manner. In rodents, early-life caffeine exposure induces sex-specific alterations in hippocampal glutamatergic and GABAergic synaptic markers that are accompanied by increased anxiety-like behavior [4]. More broadly, developmental caffeine exposure has been associated with persistent behavioral alterations, including deficits in spatial learning and object recognition, as well as changes in exploratory and anxiety-related behaviors [5–7]. Consistent with these experimental findings, longitudinal human studies have linked prenatal caffeine exposure to adverse neurodevelopmental outcomes. In a French birth cohort, maternal caffeine consumption during pregnancy was associated with behavioral and cognitive alterations during early childhood [8], while a more recent study reported associations with structural brain changes, including reduced cortical thickness in 10-year-old children [9].

Despite accumulating evidence that developmental caffeine exposure affects brain maturation, the developmental processes that are most vulnerable remain poorly defined. Most previous studies have examined prolonged prenatal or perinatal exposure, making it difficult to identify the specific developmental stages responsible for the observed phenotypes. Synaptogenesis represents a particularly critical period because neuronal connectivity is established through the coordinated formation, maturation, and refinement of excitatory and inhibitory synapses. Adenosine receptors, particularly A_2A_R, regulate many of these developmental processes [10,11], suggesting that transient antagonism during synaptogenesis may have lasting consequences on circuit assembly.

In the present study, we therefore investigated how caffeine exposure restricted to the period of hippocampal synaptogenesis affects excitatory and inhibitory synaptic development, neuronal function, behavior, and seizure susceptibility. Given the growing evidence for sexual dimorphism in both adenosine signaling and brain maturation, we further examined whether these developmental effects differ between males and females. Our results reveal that transient caffeine exposure during synaptogenesis induces sex-specific and temporally distinct remodeling of hippocampal circuits, leading to divergent behavioral and neuronal excitability outcomes.

## Materials and Methods

### Animals

All experimental procedures were conducted in accordance with European Directive 2010/63/EU and approved by the French Ministry of Higher Education and Research (APAFIS#8990 / 20170222104585_v4; APAFIS#38106 / 2022072811529284_v5), following evaluation by the local ethics committee CEEACD 005. All efforts were made to minimize animal suffering and to reduce the number of animals used. C57BL/6JRj mice, supplied by Janvier Lab, were delivered to our animal facility at least a week before starting the experiments. Animals were housed in standard laboratory cages on a 12-hours light/dark cycle, in a temperature-controlled room (21°C) with free access to food and water.

For chronic neonatal treatment, C57BL/6 wild-type littermates received daily intraperitoneal injections of saline or caffeine (1 mg/kg) from P3 to P16. This dose produces peak blood caffeine concentrations of approximately 1 mg/L, assuming that body weight approximates body fluid volume, which is within the range reported in umbilical cord blood (0.5–2 mg/L; [12]) and breast milk (0.32–1.15 mg/L; [13]) from caffeine-consuming mothers.

Animals remained with their mother in their home cage between injections. Behavioural tasks were performed at P25-35 or P65-80 and brains were collected for immunohistochemistry or electrophysiology at P20 or P75. Prior to perfusion, mice were anesthetized with an intraperitoneal injection of ketamine (150 mg/kg) and xylazine (15 mg/kg) diluted in a NaCl solution.

### Immunohistochemistry

To quantify VGlut1 and VGAT upon chronic caffeine treatment*, w*ild-type C57Bl6 (Janvier Labs) mice received daily intraperitoneal injections of ether saline or caffeine (1 mg/kg) between P3 and P16. Animals were left with their mother in the cage of origin during the injections. P20-21 or P75 mice were then perfused in the heart with 4% PFA in 0.12 M phosphate buffer (PB) and brains were post-fixed overnight at 4°C. Fixed brains were cryoprotected in 30% saccharose in PB and 50 µm-floating coronal sections were cut with a cryotome before immunohistochemistry. Standard immunohistochemistry was performed on free-floating sections. Slices were permeabilized in 0.5% Triton X-100 and blocked using 10% goat serum for 4 hours at room temperature (RT) and subsequently incubated in guinea pig anti-VGlut1 (1:1000, Millipore, AB5905) and rabbit anti-VGAT (1:500, Synaptic System) for 48 hours at 4°C. Slices were rinsed 3x in 0.1M PB and incubated in donkey-anti guinea pig Cy3-conjugated (1:500, Jackson Immunoresearch, 706-605-152) and donkey-anti rabbit A647 secondary antibody (1:500, Jackson Immunoresearch, 711-605-152). Images were acquired on either a Leica TCS SP5 upright confocal microscope or a Leica SP8 confocal microscope, both equipped with a 63x oil-immersion objective and operated with LAS X software (Leica). Series of z-stacks of 10 µm were acquired using a z spacing of 0.3 µm. Images were filtered on Metamorph (background and shading correction) after a Z-stack maximum projection of the signal. Then using ImageJ software, VGlut1 and VGAT cluster number were quantified in every layer of the CA3, CA1 and DG regions of the hippocampus with the “Find maxima” tool. The cluster number was obtained in 1 to 3 different squares of 150 *150 pixels in each layer when it was possible.

### mIPSC and mEPSC recordings in hippocampal slices

To analyse physiological properties of excitatory synapses after chronic treatment with caffeine during synaptogenesis *in vivo*, miniature inhibitory postsynaptic currents (mIPSCs) and miniature excitatory postsynaptic currents (mEPSCs) were recorded from slices of P20 mice. Animals were deeply anesthetized with ketamine/xylazine (115/15 mg/kg) and transcardially perfused with an ice-cold choline-based solution containing (in mM): 110 Choline Cl, 25 Glucose, 25 NaHCO_3_, 11.6 Ascorbic acid, 3.1 Pyruvic acid, 1.25 NaH_2_PO_4_, 2.5 KCl, 0.5 CaCl_2_, 7 MgCl_2_ saturated with 95% O_2_ / 5% CO_2_. Mice were then decapitated, hippocampi were rapidly dissected and 400mm transverse sections were prepared using a vibratome (Microm, Thermofisher). Slices were then transferred and allowed to recover for 1 hour in a humidified interface chamber filled with bicarbonate-buffered ACSF preheated at 37C and oxygenated with 5%CO2inO2, containing (in mM): 126 NaCl, 26 NaHCO_3_,10 Glucose, 3.5 KCl, 1.25 NaH_2_PO_4_, 1.6 CaCl_2_, 1.2 MgCl_2_. For recordings, slices were transferred in a submerged recording chamber and superfused with ACSF maintained at 32C. Neurons in CA3 region were recorded at 31°C under superfusion with a HEPES-buffered, artificial CSF (H-ACSF) containing (in mM) the following : 125 NaCl, 20 D-glucose, 10 HEPES, 4 MgCl_2_, 2 KCl, 1 CaCl_2_, pH7.4. Cells were held at +10mV for mIPSC recording, while miniature excitatory postsynaptic currents (mEPSCs) were recorded at a holding potential of −70 mV. mIPSCs were isolated by adding TTX (1 µM) and mEPSC by adding TTX (1 µM) and bicuculline methochloride (20 µM) to the extracellular solution. Currents were recorded with a Multiclamp 700B amplifier (Molecular Devices), filtered at 2 kHz, and digitized at 20 kHz. Access and input resistance were regularly monitored with 5mV voltage steps. mIPSC and mEPSCs were detected and analyzed offline using Detectivent software [14].

### Behavior

#### Open field

Behavioral testing was performed on mice between 9:00 am - 3:00 pm. The test took place in a white square open-field (50 x 50 cm for P30 and 1 x 1 m for P75 mice) apparatus under high illumination (50 lux for P30 and 300 lux for P75 mice). The central area of the open field was a square of 25% of the area. Animals were placed in a corner and were allowed to freely explore the open-field. A video tracking system, which included a computer-linked overhead camera allowing gravity center detection, was used to monitor general locomotor activity and activity in the center every min for 9 consecutive minutes (ViewPoint, Lyon, France). Locomotor activity data were collected from different cohorts. To account for differences between control groups, data were normalized to the mean value of the respective control group, allowing comparison across cohorts.

#### Spatial memory

Mice were tested successively for the detection of novel spatial position of an object and of a novel visual object. The test took place in a square arena (50 x 50 cm) apparatus under low illumination (< 30 lux for P30 and < 50 lux for P75 mice). On the testing day, mice were placed in the arena and were allowed to explore two identical objects during 4 acquisition trials of 5 minutes each and separated by a 3 minutes inter-trial interval. On the fifth trial, mice were placed back into the arena in which one of the objects was moved to a new spatial position (place recognition).

A video tracking system, which included a computer-linked overhead camera allowing nose detection, was used to monitor exploration of each wire pot every minute of each trial (ViewPoint, Lyon, France). A preference index was used as a measure of discrimination between novel and old locations was obtained as follows :

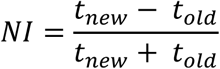

Automatic animal tracking was completed using the ViewPoint analysis software.

#### Social exploration

The test took place in a square arena (50 x 50 cm) apparatus under low illumination (< 50 lux). Mice were first allowed to habituate to the arena for 10 minutes, then habituated to the chamber containing the empty small round wire pot for 10 minutes. For the final stage, an unfamiliar mouse was placed under one wire pot and an object under the other one. Mice were left to freely explore for 10 minutes. A video tracking system, which included a computer-linked overhead camera allowing nose detection, was used to monitor exploration of each wire pot every minute for each trial (ViewPoint, Lyon, France). A preference index (PI) was used as a measure of preference for the unfamiliar mouse as compared to the object and was obtained as follows :

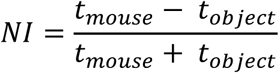

#### PTZ mice seizure

Saline or caffeine treated mice were injected subcutaneously at P20 with pentylenetatrazole (PTZ, 40 mg per kg of body weight, dissolved in saline), and recorded right after injection using video recordings. The sampling of animals as well as the experimental procedure and analysis of the data were determined based on previous published work. After 1 hour of observation, animals were sacrificed by cervical dislocation. The severity of PTZ-induced convulsive seizures was measured as in previous publications [15–17]. The scale scores from 0 (no abnormal behavior) (1) hypoactivity and postration (2) partial clonus, including that of the face, head, or forelimbs (3) generalized clonus including extremities and the tail (4) tonico-clonic seizure (5) death.

### Statistics

Experimenters were blind to the condition of the sample analyzed. Sample size selection for experiments was based on published experiments, pilot studies as well as in-house expertise. For each experiment, statistical details (n, p-values, and test used) can always be found in the figure legends. All statistical analyses were performed using GraphPad Prism 10.1.0 (Dotmatics, Boston, USA. Normally and non-normally data are presented as means ± standard error of the mean. Normality of distribution was assessed by the Shapiro-Wilk test. Normally distributed unpaired datasets were compared using the unpaired t test, ordinary one-way ANOVA tests followed by Holm-Sidak’s multiple comparison tests or two-way ANOVA tests followed by Sidak’s multiple comparison for kinetic analyses. Non-Gaussian unpaired datasets were tested by two-tailed unpaired non-parametric Mann-Whitney test or Kruskal–Wallis tests followed by Dunn’s multiple comparison tests. Cumulative distributions were compared with the Kolmogorov-Smirnov test. *P*-values < 0.05 were considered statistically significant. Indications of significance corresponding to p-values < 0.05 (*), p < 0.01 (**), p < 0.001 (***) are reported in the figures and in the corresponding legends.

## Results

### Caffeine exposure during synaptogenesis induces opposite sex-dependent temporal patterns of hippocampal glutamatergic synapse remodeling

To determine whether caffeine exposure during the critical period of synaptogenesis produces long-lasting alterations in hippocampal synaptic organization, mice were exposed to caffeine or saline during synaptogenesis i.e. between P3 and P16 and glutamatergic synapse density was analyzed at P20 and adulthood (P75). VGlut1-positive synaptic clusters were quantified across different hippocampal layers of the CA3, CA1, and dentate gyrus (DG) regions in males and females (Figure 1).

**Figure 1.**
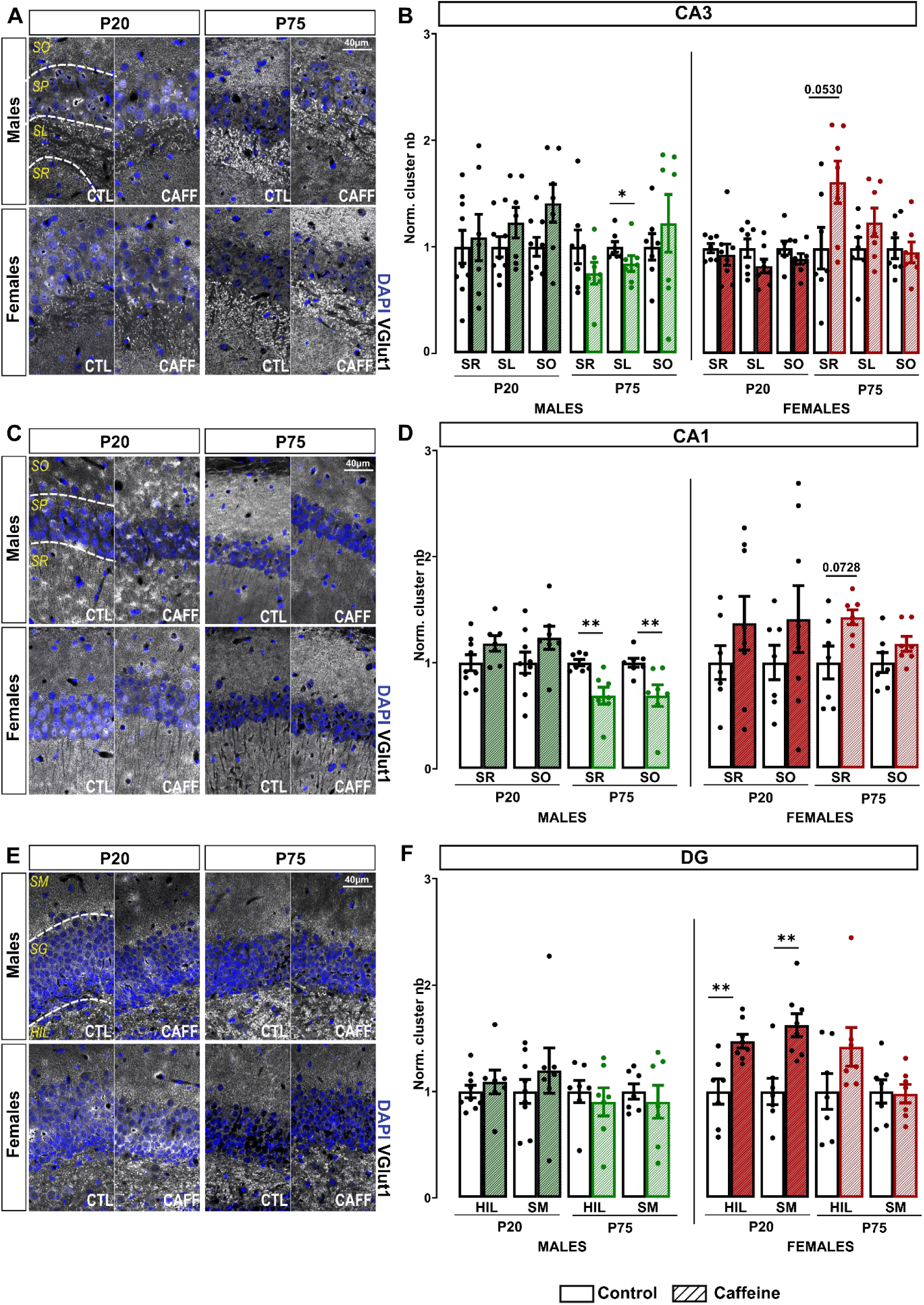
Caffeine intake during synaptogenesis leads to long-term sex-dependent effects on glutamatergic synapses in the hippocampus. **(A, C, E)** Representative images of DAPI (blue) and VGlut1 (gray) immunostaining in the stratum oriens (SO), stratum pyramidale (SP), stratum lucidum (SL), stratum radiatum (SR), hilus (HIL), and stratum moleculare (SM) of the CA3 (A), CA1 (C), and dentate gyrus (E) regions of the hippocampus from animals treated with saline (CTL) or caffeine (CAFF) during synaptogenesis. **(B, D, F)** Quantification of VGlut1 cluster density in the different layers of the CA3 (B), CA1 (D), and dentate gyrus (DG) (F) at P20 and P75 in males (left) and females (right). N_Males,P20,NaCl_ = 9, N_Males,P20,Caff_ = 6-7, N_Males,P75,NaCl_ = 7, N_Males,P75,Caff_ = 7, N_Females,P20,NaCl_ = 7, N_Females,P20,Caff_ = 7-8, N_Females,P75,NaCl_ = 7, N_Females,P75,Caff_ = 7. In all graphs, values were normalized to the corresponding control layer values; means ans s.e.m. are represented. Mann Whitney test. *p<0,05; **p<0,01 p values: **CA3 males**, P20: SR, p= 0.8371; SL, p= 0.2105; SO, p= 0.1142; P75: SR, p= 0.3829; SL, p= 0.0175; SO, p= 0.5350. **CA3 females**, P20: SR, p= 0.3357; SL, p= 0.1520; SO, p= 0.1893; P75: SR, p= 0.0530; SL, p= 0.2593; SO, p= 0.9015. **CA1 males**, P20: SR, p= 0.0907; SO, p= 0.1142; P75: SR, p= 0.0070; SO, p= 0.0023. **CA1 females**, P20: SR, p= 0.3357; SO, p= 0.3969; P75: SR, p= 0.0728; SO, p= 0.1649. **DG males**, P20: HIL, p= 0.4079; SM, p= 0.5360; P75: HIL, p= 0.4557; SM, p= 0.9015. **DG females**, P20: HIL, p= 0.0037; SM, p=0.0037; P75: HIL, p= 0.2086; SM, p= 0.9015.

In the CA3 region, caffeine exposure did not significantly modify VGlut1 cluster density at P20 in either males or females. At P75, a significant decrease in VGlut1 cluster density was observed in the stratum lucidum (SL) of caffeine-treated males compared with controls (Figure 1A–B; p = 0.0175), whereas no significant changes were detected in the other CA3 layers. In females, caffeine exposure did not significantly affect VGlut1 cluster density in CA3 at adulthood, although a tendency toward an increase was observed in the stratum radiatum (SR) (Figure 1A–B; p = 0.0530).

In the CA1 region, caffeine-induced alterations were more pronounced in males. While no significant changes were detected at P20, adult caffeine-treated males displayed a significant decrease in VGlut1 cluster density in both the stratum radiatum and stratum oriens (SO) (Figure 1C–D; SR: p = 0.0070; SO: p = 0.0023). In contrast, females showed a trend toward increased CA1 glutamatergic synapse density in the stratum radiatum (SR) at P20 and P75 (Figure 1C–D).

In the DG, caffeine exposure produced an early, transient effect in females. At P20, caffeine-treated females exhibited a significant increase in VGlut1 cluster density in both the hilus and stratum moleculare (SM) (Figure 1E–F; HIL: p = 0.0037; SM: p = 0.0037), an effect that was no longer detected at P75. No significant changes were observed in DG glutamatergic synapses in males at either developmental stage (Figure 1E–F).

Together, these findings demonstrate that caffeine exposure during synaptogenesis induces sex-dependent remodeling of hippocampal glutamatergic synapses, with a delayed reduction in excitatory synapse density in males and an earlier, transient increase in females, revealing distinct temporal trajectories as well as opposite patterns of synaptic remodeling between the two sexes.

### Developmental caffeine exposure induces a transient male-specific increase in inhibitory synaptic density in CA3

We next investigated whether caffeine exposure affected inhibitory synapse formation by quantifying VGAT-positive clusters in hippocampal regions at P20 and P75 (Figure 2). In the CA3 region, caffeine treatment induced an increase in inhibitory synaptic density during development in males. Specifically, caffeine-treated males displayed increased VGAT cluster density in the stratum lucidum and stratum oriens at P20 compared with saline controls (Figure 2A–B; SL: p = 0.0381; SO: p = 0.0095). These changes were no longer observed at P75, suggesting a transient developmental effect. In females, caffeine exposure did not significantly modify VGAT cluster density in CA3 at either P20 or P75 (Figure 2A–B). No major alterations in inhibitory synaptic density were observed in the CA1 (Figure 2C–D) or DG (Figure 2E–F) regions in either sex,

**Figure 2.**
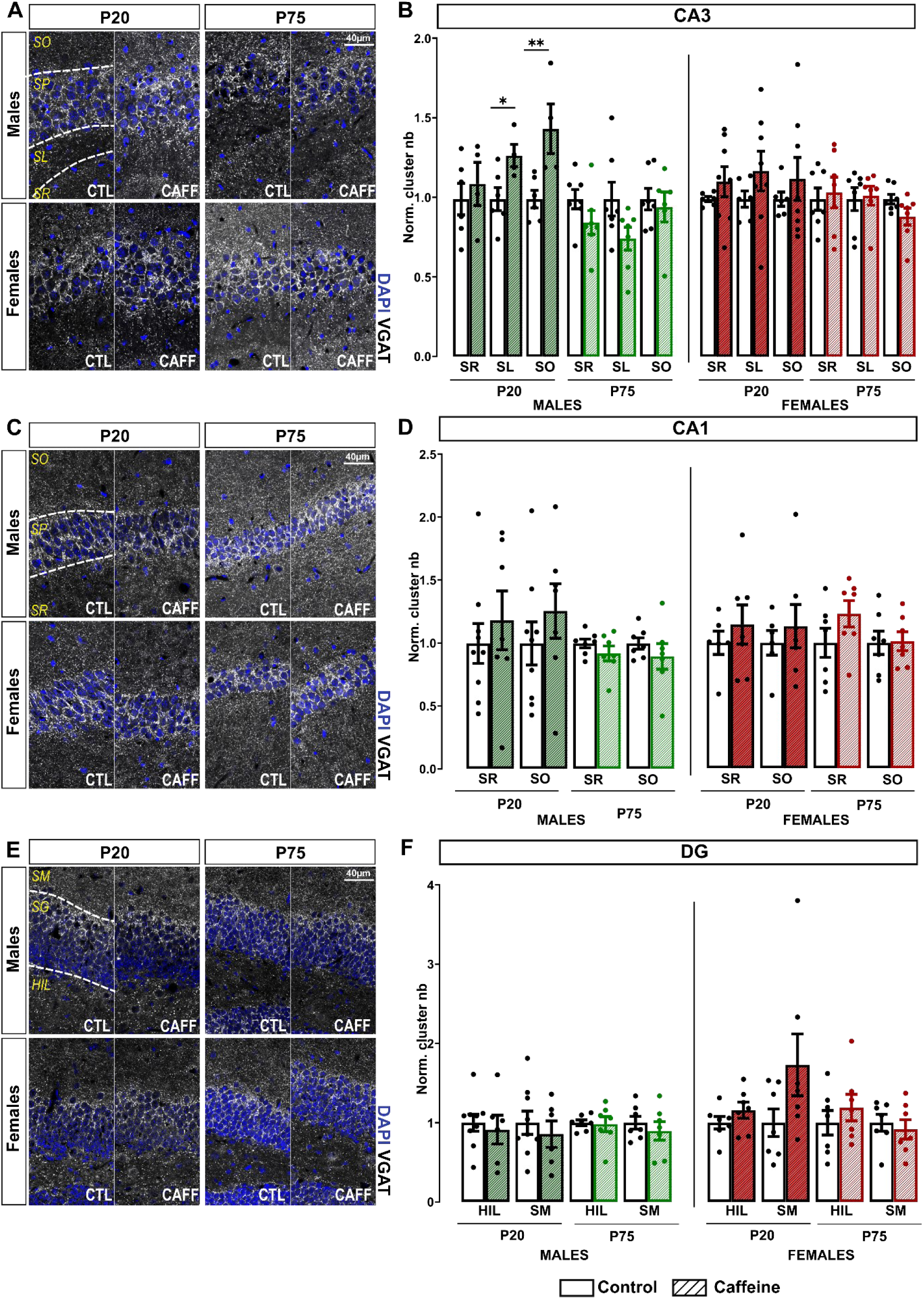
Caffeine intake during synaptogenesis leads to a transient increase in inhibitory synapses in the CA3 of males. **(A, C, E)** Representative images of DAPI (blue) and VGAT (gray) immunostaining in the stratum oriens (SO), stratum pyramidale (SP), stratum lucidum (SL), stratum radiatum (SR), hilus (H), and stratum moleculare (SM) of the CA3 (A), CA1 (C), and dentate gyrus (E) regions of the hippocampus from animals treated with saline (CTL) or caffeine (CAFF) during synaptogenesis. **(B, D, F)** Quantification of VGAT cluster density in the different layers of the CA3 (B), CA1 (D), and dentate gyrus (DG) (F) at P20 and P75 in males (left) and females (right). N_Males,P20,NaCl_ = 6-9, N_Males,P20,Caff_ = 4-7, N_Males,P75,NaCl_ = 7, N_Males,P75,Caff_ = 7, N_Females,P20,NaCl_ = 5-7, N_Females,P20,Caff_ = 5-8, N_Females,P75,NaCl_ =6-7, N_Females,P75,Caff_ = 7. In all graphs, values were normalized to the corresponding control layer values; means ans s.e.m. are represented. Mann Whitney test. *p<0,05. P values: **CA3 males**, P20: SR, p= 0.6095; SL, p= 0.0381; SO, p= 0.0095; P75: SR, p= 0.0973; SL, p= 0.0973; SO, p= 0.9452. **CA3 females**, P20: SR, p= 0.4908; SL, p= 0.2284; SO, p>0.9999; P75: SR, p= 0.8048; SL, p= 0.9015; SO, p= 0.2086. **CA1 males**, P20: SR, p= 0.7577; SO, p= 0.3510; P75: SR, p= 0.4557; SO, p= 0.3829. **CA1 females**, P20: SR, p= 0.6282; SO, p=0.8357; P75: SR, p= 0.2739; SO, p=0.9015. **DG males**, P20: HIL, p= 0.4559; SM, p= 0.6070; P75: HIL, p= 0.8048; SM, p= 0.7401. **DG females**, P20: HIL, p= 0.2593; SM, p= 0.2086; P75: HIL, p= 0.3829; SM, p= 0.6200.

Overall, caffeine exposure during synaptogenesis preferentially affects inhibitory synapse organization in the CA3 region in males, inducing a transient increase during development with limited long-term consequences, whereas females show no significant alterations.

### Developmental caffeine exposure modifies synaptic transmission in a sex-dependent manner

To determine whether caffeine-induced changes in synaptic density were associated with functional alterations, spontaneous miniature excitatory and inhibitory postsynaptic currents (mEPSCs and mIPSCs) were recorded from CA3 pyramidal neurons at P20 (Figure 3).

**Figure 3.**
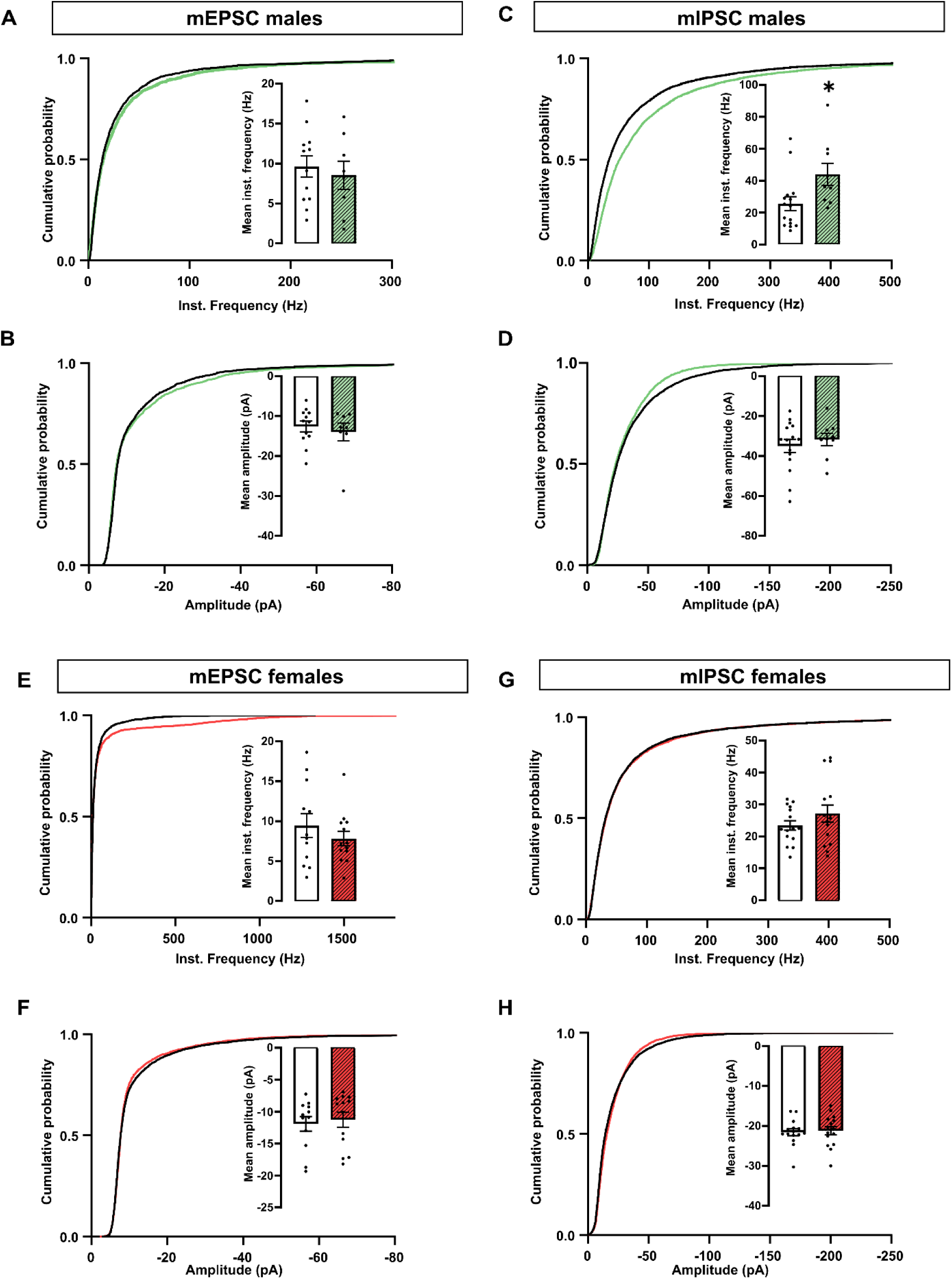
Caffeine intake during synaptogenesis increases mIPSC frequency in males and tends to increase mEPSC frequency in females. **(A, B)** Quantification of mEPSC amplitude (A) and frequency (B) in CA3 pyramidal neurons from P20 males treated with saline (white) or caffeine (green). **(C, D)** Quantification of mIPSC amplitude (C) and frequency (D) in CA3 pyramidal neurons from P20 males treated with saline (white) or caffeine (green). **(E, F)** Quantification of mEPSC amplitude (E) and frequency (F) in CA3 pyramidal neurons from P20 females treated with saline (white) or caffeine (green). **(G, H)** Quantification of mIPSC amplitude (G) and frequency (H) in CA3 pyramidal neurons from P20 females treated with saline (white) or caffeine (green). n_mEPSC_=12 cells (saline, males), n_mEPSC_=10 (caffeine, males), n_mIPSC_=15 cells (saline, males), n_mIPSC_=8 cells (caffeine, males), n_mEPSC_=11 cells (saline, females), n_mEPSC_=12 (caffeine, females), n_mIPSC_=15 cells (saline, females), n_mIPSC_= 15 cells (caffeine, females). For histograms means and s.e.m. are represented; Welch’s t-test for normally distributed data and Mann-Whitney for non-normally distributed data. *p < 0.05 P-value : **mEPSC males**, Frequency, p= 0.6296; Amplitude, p= 0.7345. **mIPSC males**, Frequency, p= 0.0212; Amplitude, p= 0.4859. **mEPSC females**, Frequency, p= 0.3590; Amplitude, p= 0.3475. **mIPSC females**, Frequency, p= 0.2332; Amplitude, p= 0.6236.

In males, caffeine exposure did not significantly modify mEPSC frequency or amplitude (Figure 3A–B). In contrast, caffeine-treated males displayed a significant increase in mIPSC frequency compared with saline controls, while mIPSC amplitude remained unchanged (Figure 3C–D), indicating an increase in inhibitory synaptic input without detectable changes in postsynaptic receptor function. This is consistent with the selective increase in VGAT-positive inhibitory synapses observed by immunohistochemistry, while VGlut1-positive excitatory synapses remained unchanged at P20 (Figure 2A–B).

In females, caffeine exposure did not significantly alter mEPSC (Figure 3E–F) or mIPSC (Figure 3G–H) amplitude or frequency, in agreement with the lack of detectable changes in VGlut1- and VGAT-positive synaptic density observed at P20 in CA3 (Figure 2B, 3B).

These findings indicate that caffeine exposure during synaptogenesis produces sex-specific functional consequences, enhancing inhibitory transmission in males during early development.

### Caffeine exposure during synaptogenesis induces sex-dependent behavioral alterations

We next investigated whether caffeine exposure during synaptogenesis affected behavioral outcomes during adolescence and adulthood. Developmental caffeine exposure did not alter locomotor activity in either males or females at P30 or P75, as indicated by the total distance traveled in the open field test (Figure S1A–B).

The same test was used to assess anxiety-like behavior. Caffeine exposure during synaptogenesis induced an anxiety-like phenotype selectively in males. Adolescent and adult caffeine-treated males exhibited reduced exploration of the center of the open field, as reflected by decreased active exploration distance and duration in center compared with control animals (Figure 4A–C). In contrast, caffeine exposure did not significantly alter open-field exploration in females at either P30 or P75 (Figure 4D–F).

**Figure 4.**
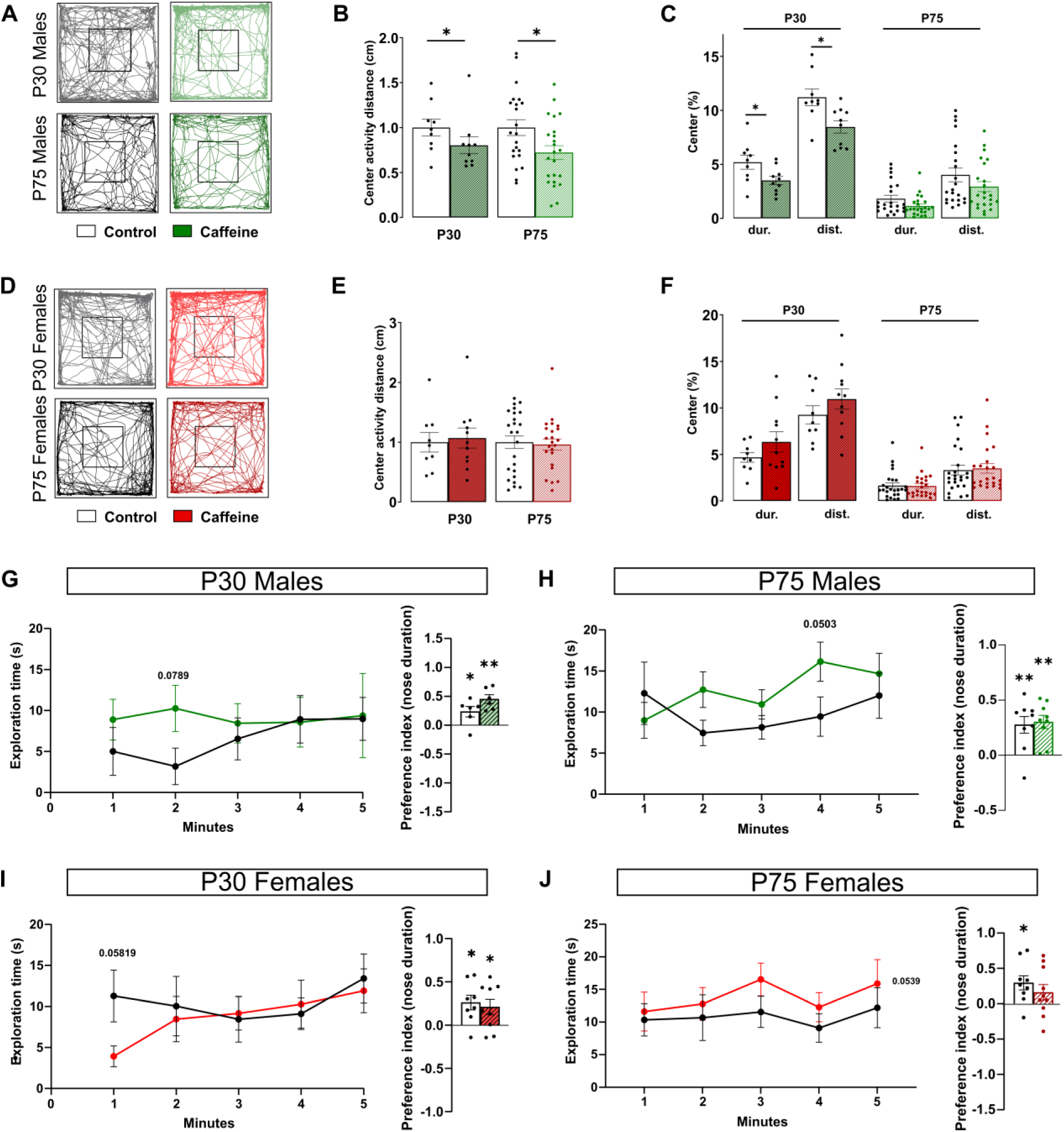
Caffeine intake during synaptogenesis induces anxiety-like behavior in males and spatial recognition memory deficits in adult females. **(A, D)** Representative open field (OF) tracks from P30/P75 males (A) and females (D) treated with saline (CTL) or caffeine (CAFF) during synaptogenesis. **(B, E)** Quantification of total active exploration in the center of the OF in males (B) and females (E). **(C, F)** Percentage of active exploration duration and distance in the center relative to total active exploration in the OF in males (C) and females (F). P75 mice correspond to the mice tested at P30 and followed until adulthood. At P30: N_Males,NaCl_ = 9, N_Males,Caff_ = 10, N_Females,NaCl_ = 8, N_Females,P75,Caff_ = 11; at P75 N_Males,NaCl_ = 22, N_Males,Caff_ = 23, N_Females,NaCl_ = 25, N_Females,P75,Caff_ = 23. Values were normalized to the corresponding control values; data are presented as mean ± s.e.m. Welch’s t-test for normally distributed data and Mann-Whitney for non-normally distributed data. *p < 0.05, **p < 0.01. P-value **P30 Males**, Center activity distance, p= 0.0191; Active center exploration duration (%), p= 0.0416; Active center exploration distance (%), p= 0.0113. **P75 Males**, Center activity distance, p= 0.0217; Active center exploration duration (%), p= 0.0989; Active center exploration distance (%), p= 0.1712. **P30 Females**, Center activity distance, p= 0.8238; Active center exploration duration (%), p= 0.1929; Active center exploration distance (%), p= 0.2620. **P75 Females**, Center activity distance, p= 0.5125; Active center exploration duration (%), p= 0.9468; Active center exploration distance (%), p= 0.9348. **(G–J)** Quantification of total exploration over time and place recognition test performance (histograms) in P30 males (G), P75 males (H), P30 females (I), and P75 females (J) treated with saline or caffeine during synaptogenesis. Values were normalized to the corresponding control values; data are presented as mean ± s.e.m. 2way ANOVA and one sample t-test weres used for statistical comparisons. *p < 0.05, **p < 0.01. P-value: **P30 Males**, Exploration time, Time x treatment p= 0.6038, Time p= 0.7648, Treatment p= 0.2551. T-test control, p=0.0496, T-test caffeine, p= 0.0018. **P75 Males**, Exploration time, Time x treatment p= 0.2096, Time p= 0.3278, Treatment p= 0.1560. T-test control, p=0.0065, T-test caffeine, p= 0.0010. **P30 Females**, Exploration time, Time x treatment p= 0.4803, Time p= 0.3849, Treatment p= 0.3736. T-test control, p= 0.0170, T-test caffeine, p= 0.0440. **P75 Females**, Exploration time, Time x treatment p= 0.9464, Time p= 0.5965, Treatment p= 0.0539. T-test control, p=0.0178, T-test caffeine, p= 0.1801.

Spatial memory was then assessed using the novel object location (NOL) tasks. In contrast to the anxiety-like phenotype observed exclusively in males, developmental caffeine exposure selectively impaired recognition memory in females. No difference regarding the distance during the task was observed (Figure S1C-D), but adult caffeine-exposed females exhibited reduced performance in the novel object location task (Figure 4I-J), whereas males showed no significant deficits in NOL (Figure 4G–H) test at either developmental stage. Notably, the recognition memory deficit in caffeine-exposed females emerged only at P75 and was absent at P30, indicating a delayed manifestation of the cognitive consequences of developmental caffeine exposure.

Social preference was assessed in adulthood using the nose exploration test, in which mice were allowed to explore either a novel conspecific or a non-social object for 10 min (Figure S2A). Developmental caffeine exposure did not alter social preference in either males or females (Figure S2B), indicating that transient caffeine exposure during synaptogenesis does not impair social preference in adulthood.

Together, these findings reveal distinct long-term, sex-dependent behavioral outcomes following developmental caffeine exposure, characterized by increased anxiety-like behavior in males, and selective recognition memory impairment in females.

### Developmental caffeine exposure selectively increases PTZ-induced seizure susceptibility in adolescent females

Finally, we investigated whether caffeine exposure during synaptogenesis altered seizure susceptibility using the PTZ-induced seizure model at P20 and P75 (Figure 5). In males, caffeine exposure did not significantly affect seizure progression, maximum Racine score, time spent at each seizure stage, or the proportion of animals reaching stage 5 during the first hour following PTZ injection, either at P20 (Figure 5A–D) or at P75 (Figure 5I–L).

**Figure 5.**
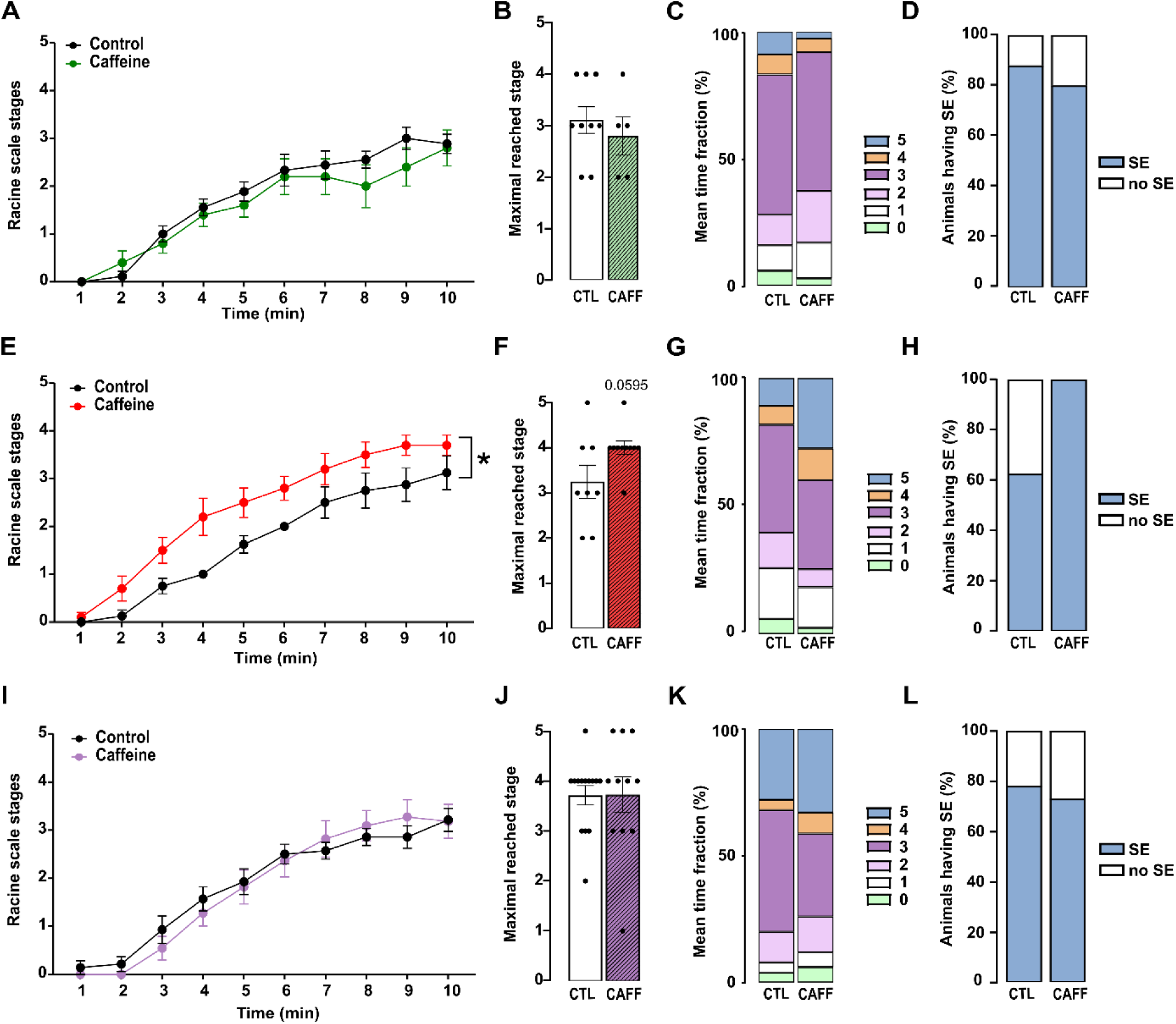
Caffeine intake during synaptogenesis increases PTZ seizure susceptibility in P20 females. **(A, E, I)** Mean Racine score per minute during the first 10 min after PTZ injection in P20 males (A), P20 females (E) and P75 mice (I) treated with saline (CTL) or caffeine (CAFF) during synaptogenesis. **(B, F, J)** Maximum Racine stage reached during the first 10 min after PTZ injection in P20 males (B), P20 females (F) and P75 mice (J) treated with saline (CTL) or caffeine (CAFF). **(C, G, K)** Mean fraction of time spent at each Racine stage per group (%) in P20 males (C), 20 females (G) and P75 mice (K) during the 1 h following PTZ injection. **(D, H, L)** Percentage of animals reaching Racine stage 4 in P20 males (D),P20 females (H) and adult mice (L). **P20** N_Males,NaCl_ = 9, N_Males,Caff_ = 5, N_Females,NaCl_ = 8, N_Females,P75,Caff_ = 10; **P75** N_NaCl_ = 14, N_Caff_ = 11. Data are presented as mean ± s.e.m. Two-way ANOVA followed by Mann–Whitney test. *p < 0.05. P-values: **P20 males**, Racine scale time, treatment, p= 0.4093; Maximal reached stage, p= 0.7293; Mean time fraction, p= 0.9994; Animal having seizure, p>0.9999. **P20 females**, Racine scale time, treatment, p= 0.0151; Maximal reached stage, p= 0.0595; Mean time fraction, p>0.9999; Animal having seizure, p>0.9999. **Adult mice**, Racine scale time, treatment, p= 0.8628; Maximal reached stage, p= 0.7889; Mean time fraction, p>0.9999; Animal having seizure, p= 0.9976.

In contrast, caffeine-treated females exhibited increased seizure severity following PTZ administration at P20, as evidenced by higher Racine scores and an increased maximal seizure stage compared with control females (Figure 5E-H). However, this increased seizure susceptibility was absent at P75 (Figure 5I-L), demonstrating that the effect was restricted to adolescence and did not persist into adulthood.

Together, these findings reveal that caffeine exposure during synaptogenesis produces persistent and sex-dependent modifications of hippocampal synaptic organization and function, associated with distinct behavioral and seizure-related phenotypes.

## Discussion

Our findings extend previous observations by Teixeira-Silva et al.[4], who reported sex-dependent alterations in hippocampal glutamatergic and GABAergic markers following developmental caffeine exposure. However, their analyses were restricted to juvenile animals (up to P26) and relied on Western blot quantification of synaptic proteins, including NMDA and AMPA receptor subunits and GAD, providing information on overall protein abundance but not on synaptic density, regional specificity, or the functional consequences at individual synapses. In contrast, our study directly quantified excitatory and inhibitory synapses in distinct hippocampal subregions, combined these structural analyses with electrophysiological assessments, and investigated both the short- and long-term consequences of caffeine exposure during synaptogenesis. In females, Teixeira-Silva et al. reported increased GluA1 expression together with reduced GAD levels, suggesting an early shift toward enhanced excitatory drive. Consistent with this interpretation, we observed a transient increase in glutamatergic synapse density in the dentate gyrus at P20, followed by a delayed increase in excitatory synapses in the CA3 and CA1 regions that persisted into adulthood, whereas inhibitory synapse density remained largely unchanged. These findings indicate that developmental caffeine exposure induces a sustained increase in the excitatory/inhibitory (E/I) balance in females through temporally and regionally distinct patterns of synaptic remodeling. In males, Teixeira-Silva et al. described increased GluN1/GluN2B expression, reduced GAD levels, and decreased GluA1 expression, suggesting a more complex early reorganization of excitatory and inhibitory signaling. Our analyses revealed a reduction in excitatory synapse density in adulthood, particularly in the CA1 region, with no significant effect on inhibitory synapses, suggesting a long-term decrease in the E/I balance. In the CA3 region, we observed a transient increase in inhibitory synapse density without a significant change in excitatory synapses, indicating an early reduction in the E/I balance. Therefore, our results suggest that, in males, developmental caffeine exposure decreases the E/I balance both transiently during juvenile development (CA3) and persistently in adulthood (CA1).

Together, these findings demonstrate that developmental caffeine exposure induces sex-specific trajectories of hippocampal circuit remodeling that evolve throughout postnatal maturation and persist into adulthood, emphasizing that the long-term consequences of developmental caffeine exposure cannot be fully captured by early measurements of synaptic protein abundance alone. Furthermore, our data reveal opposite effects of developmental caffeine exposure on hippocampal E/I balance in males and females, with females exhibiting an increased E/I balance and males showing a reduced E/I balance.

The increased anxiety-like behavior observed in males may be related to the long-term disruption of hippocampal E/I balance. The transient increase in inhibitory synapse density in CA3 during development could alter hippocampal network dynamics by reducing circuit excitability and impairing the maturation of emotional circuits. In adulthood, the reduction of excitatory synapses in CA1, a major hippocampal output region involved in communication with limbic and prefrontal areas, may further affect the regulation of anxiety-related responses. Together, these alterations suggest that developmental caffeine exposure may impair the maturation of hippocampal circuits involved in emotional regulation.

The increased E/I balance observed in females suggests that developmental caffeine exposure may shift hippocampal circuits toward a more excitable state in females compared with males. This interpretation is consistent with our behavioral findings showing that females exhibit greater susceptibility than males to acute PTZ-induced seizures. Interestingly, the increase in excitatory synapse density observed in females dentate gyrus was transient, being significant in the dentate gyrus at P30 but no longer detectable in adulthood (P75). Likewise, the enhanced susceptibility to PTZ-induced seizures was also transient and restricted to juvenile animals. The parallel temporal profile of these structural and behavioral alterations suggests that the transient increase in excitatory synapse density within the dentate gyrus in females may contribute to the heightened seizure susceptibility observed during early postnatal development.

In females, the shift toward an increased excitatory/inhibitory balance may alter the fine regulation of hippocampal network activity required for efficient memory encoding. Indeed, excessive glutamatergic signaling may reduce neuronal signal-to-noise ratio, disrupt the balance between synaptic potentiation and depression (LTP/LTD), and impair the ability to discriminate between familiar and novel object locations. Thus, the impaired location memory observed in females at P75 may result from a persistent increase in hippocampal excitatory drive that disrupts the synaptic plasticity mechanisms required for optimal memory processing. In contrast, males exhibited a different trajectory of synaptic remodeling, characterized a tendency toward reduced hippocampal E/I balance rather than hyperexcitability. Such alterations may preferentially affect emotional regulation circuits rather than recognition memory processes, consistent with the anxiety-like phenotype observed in males.

These sex-specific outcomes may also involve the influence of sex hormones during adulthood, as estrogen signaling strongly regulates dendritic spine density, glutamatergic transmission, and hippocampal plasticity [18]. Consequently, caffeine exposure during a critical period of synaptogenesis may differentially program the maturation of hippocampal circuits in males and females, leading to distinct long-term behavioral vulnerabilities.

## Data availability statement

The datasets generated in this study are available in the Zenodo repository under accession number xxxxx.

## Acknowledgments

We thank the animal and imaging facilities of IFM and the service unit IPGG Technological Platform CNRS UAR 3750 for technical support.

## Author Contributions

S.L., S.R., M.N.B. and D.B. designed the experiments. S.R. prepared the figures and S.L. and S.R. wrote the paper. S.R. performed immunohistochemistry experiments, S.R. and Z.I. analyzed the data. J.P. performed ectrophysiological experiments and analyzed the data. S.R. conducted behavior experiments and analyzed the data. J.P. and F.G-C. performed PTZ seizure experiments and J.P., F.G-C., and M.R. analyzed the data.

## Supplementary Information

**Figure S1.**
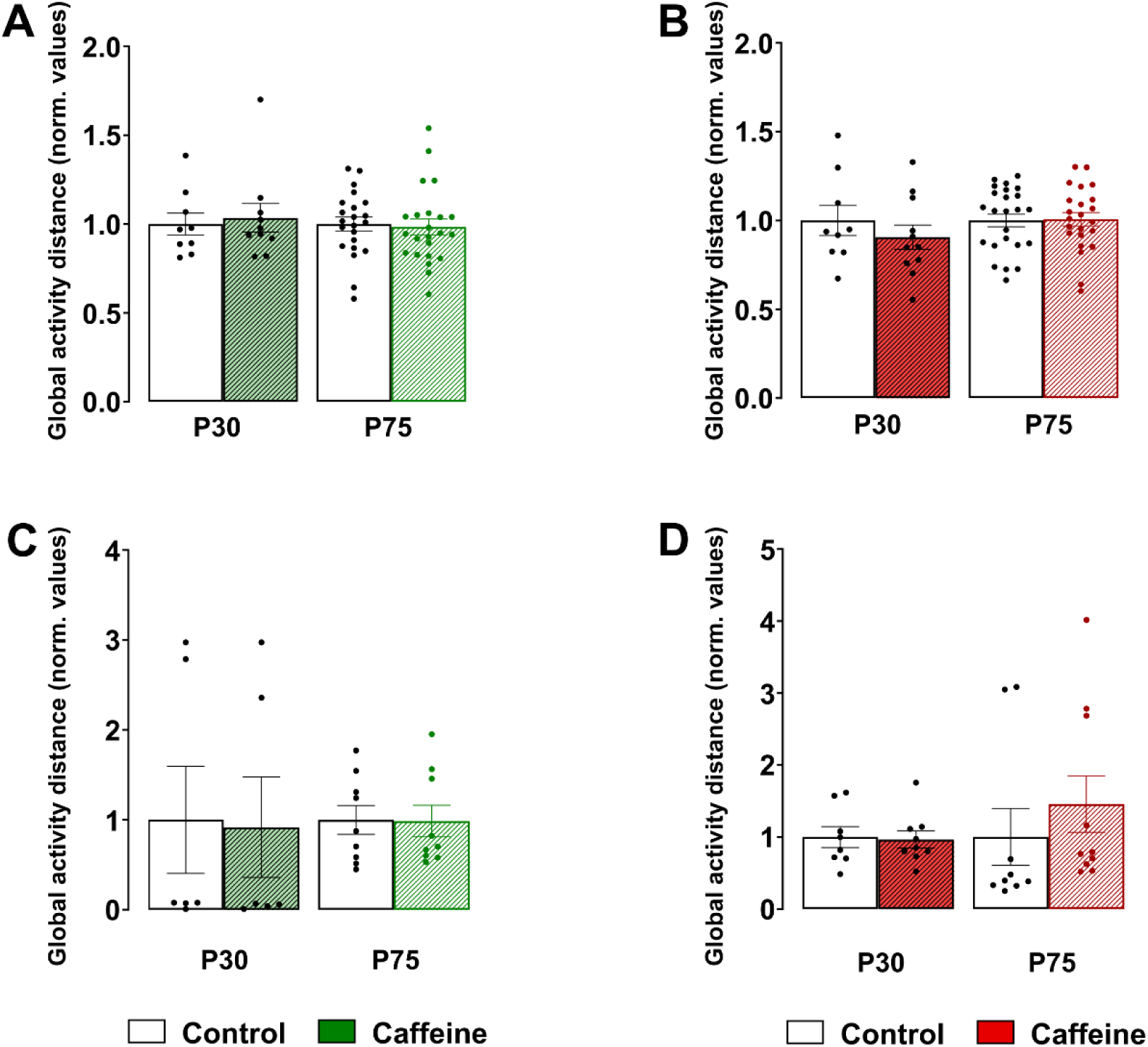
Caffeine exposure during synaptogenesis does not affect locomotor activity. **(A, B)** Normalized total distance traveled during the open field (OF) test in P30 and P75 males (A) and females (B). At P30: N_Males,NaCl_ = 9, N_Males,Caff_ = 10, N_Females,NaCl_ = 8, N_Females,P75,Caff_ = 11; at P75 N_Males,NaCl_ = 22, N_Males,Caff_ = 23, N_Females,NaCl_ = 25, N_Females,P75,Caff_ = 23. Values were normalized to the corresponding control values; data are presented as mean ± s.e.m. Mann-Whitney test. P-value : P30 Males, p= 0.9048. P75 Males, p= 0.3734. P30 Females, p= 0.3992. P75 Females, p>0.9999. **(C, D)** Normalized total distance traveled during the novel object location task in P30 and P75 males (C) and females (D). N_Males,P20,NaCl_ = 6, N_Males,P20,Caff_ = 6, N_Males,P75,NaCl_ = 9, N_Males,P75,Caff_ = 9, N_Females,P20,NaCl_ = 9, N_Females,P20,Caff_ = 5-8, N_Females,P75,NaCl_ = 9, N_Females,P75,Caff_ = 10. Values were normalized to the corresponding control values; data are presented as mean ± s.e.m. Welch’s t-test for normally distributed data and Mann-Whitney for non-normally distributed data were used for statistical comparisons. P-value : P30 Males, p= 0.4545. P75 Males, p=09549. P30 Females, p= 0.8884. P75 Females, p= 0.0535.

**Figure S2.**
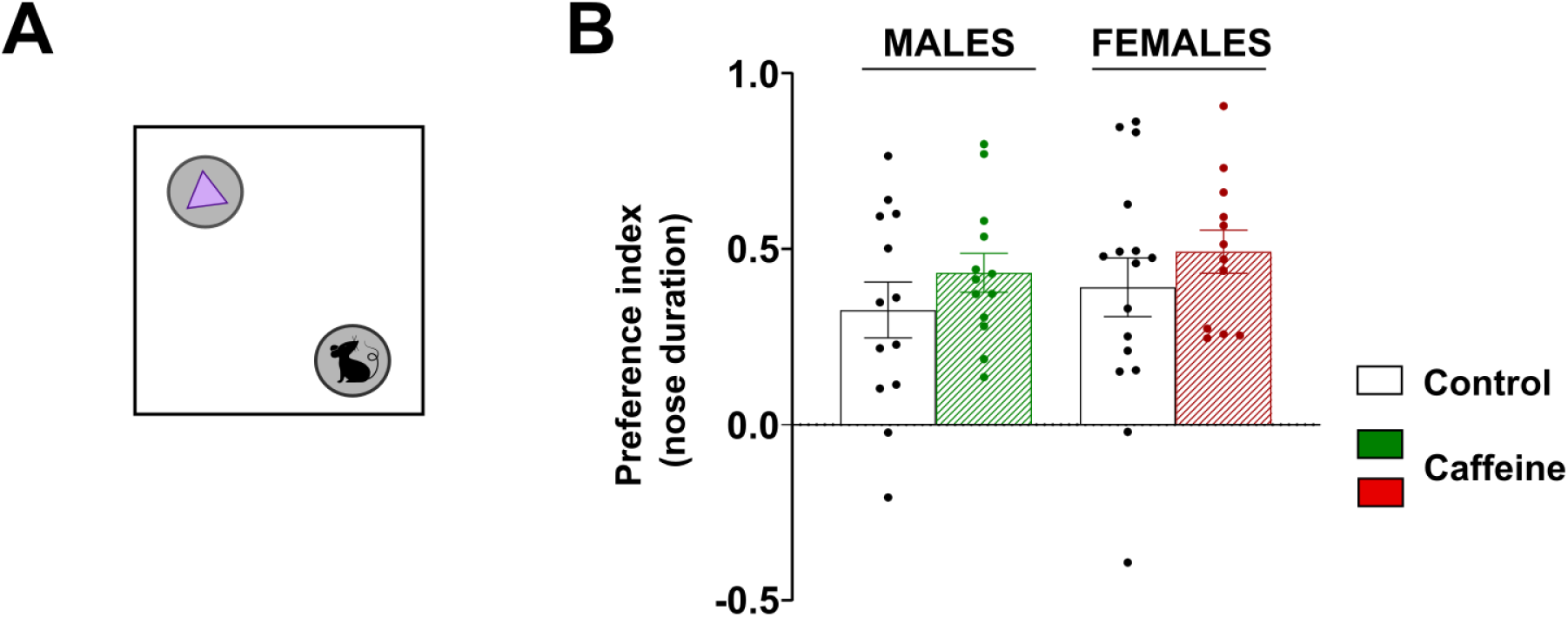
Developmental caffeine exposure does not alter social preference at P75. **(A)** Schematic representation of the social preference test performed at P75. Mice were allowed to explore either a novel conspecific or a non-social object for 10 min. **(B)** Social preference index calculated as the proportion of exploration directed toward the social stimulus. N_Males,NaCl_ = 13, N_Males,Caff_ = 13, N_Females,NaCl_ = 16, N_Females,P75,Caff_ = 12. Data are presented as mean ± s.e.m. Statistical comparisons were performed using Mann-Whitney t-tests. P values: Males p= 0.2845, Females p= 0.3658.

