## Supplementary figures and images for "Sex-specific long-term alteration of hippocampal excitation/inhibition balance and behavior by transient caffeine exposure during synaptogenesis"

### Supplemental Figure1

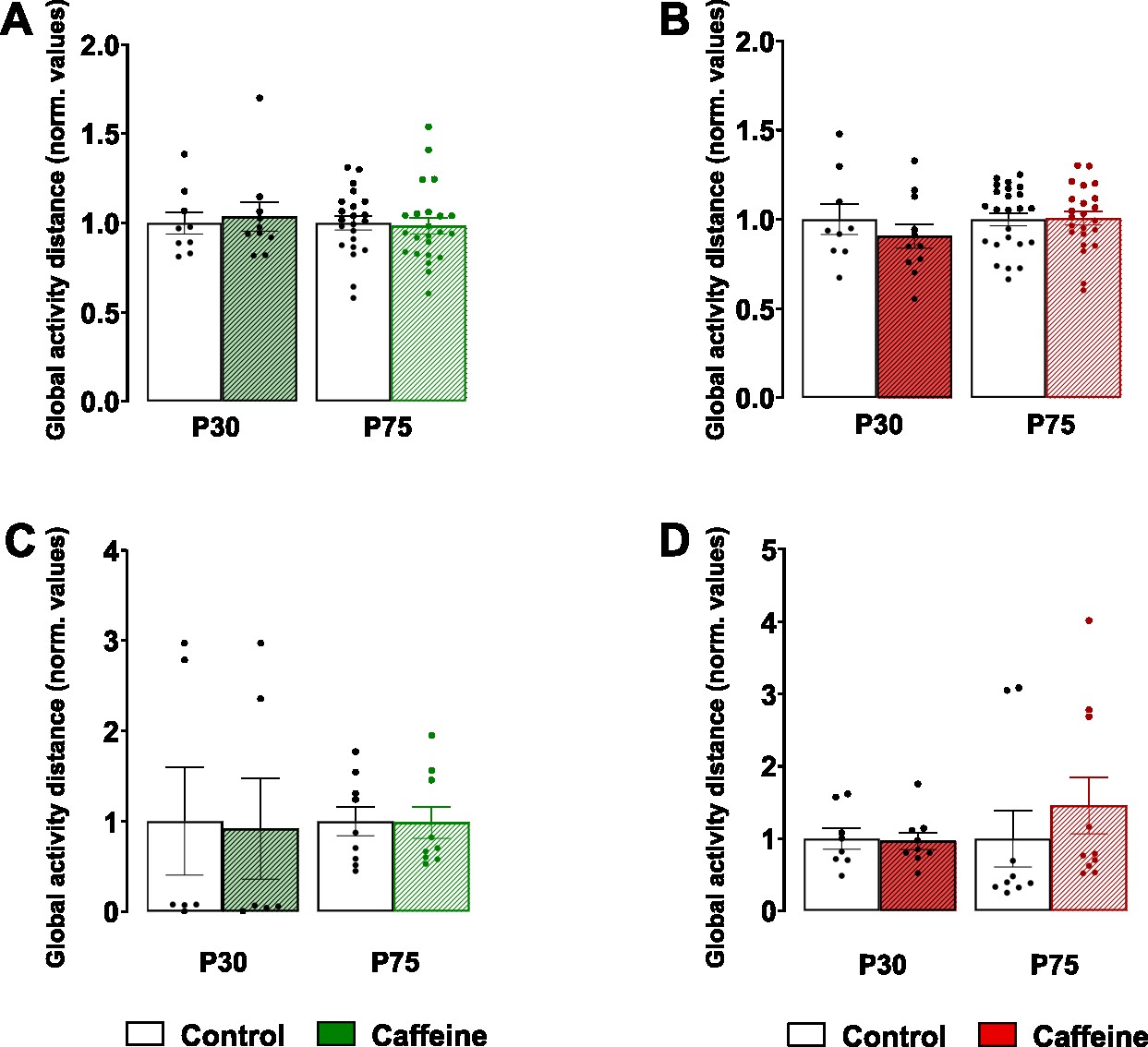

### Supplemental Figure2

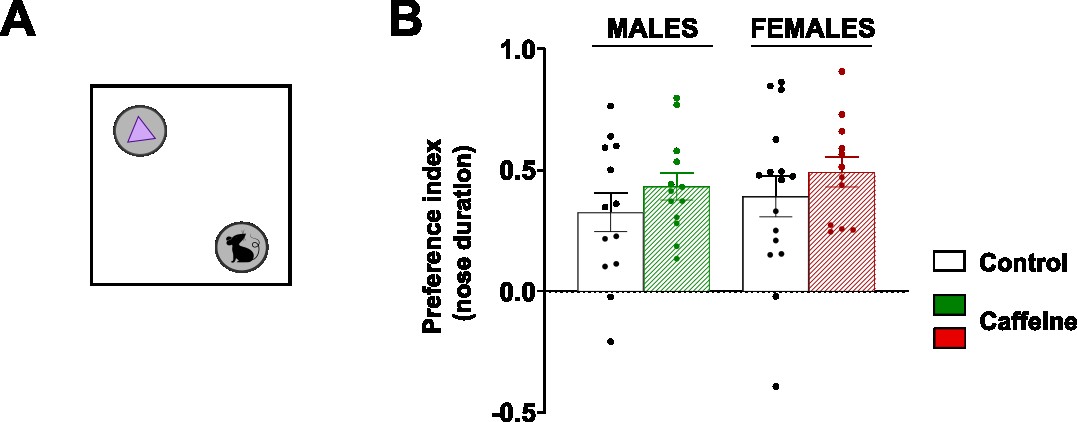
